# *effectR*: An expandable R package to predict candidate effectors

**DOI:** 10.1101/398404

**Authors:** Javier F. Tabima, Niklaus J. Grünwald

## Abstract

Effectors are by one definition small, secreted proteins that facilitate infection of host plants by all major groups of plant pathogens. Effector protein identification in oomycetes relies on identification of open reading frames with certain amino acid motifs among additional minor criteria. To date, identification of effectors relies on custom scripts to identify motifs in candidate open reading frames. Here, we developed the R package *effectR* that provides a convenient tool for rapid prediction of effectors in oomycete genomes, or with custom scripts for any genome, in a reproducible way. The *effectR* package relies on a combination of regular expressions statements and hidden Markov model approaches to predict candidate RxLR and CRN effectors. Other custom motifs for novel effectors can easily be implemented and added to package updates. The *effectR* package has been validated with published oomycete genomes. This package provides a convenient tool for reproducible identification of candidate effectors in oomycete genomes.

---

Secreted effector proteins have been reported for all major groups of plant pathogens, including bacteria, fungi, oomycetes, nematodes, and viruses (Jones and Dangl 2006). Effector proteins are defined as secreted proteins that manipulate plant processes to the advantage of the parasite, in order to promote infection and generate disease (Petre and Kamoun, 2014). Effector proteins modulate and interfere with the normal physiology of the plant host in order to facilitate disease and infection (Jones and Dangl 2006; Kamoun 2007). Notable effector proteins are common in bacterial plant pathogens. Up to 29 effector proteins are injected by the bacterial pathogen *Pseudomonas syringae* into the host plant cells via the type-3 secretion system (Chang et al. 2005). Effector discovery has predicted over 50 candidate effector proteins for the species *Heterodera glycines*, a plant-parasitic nematode that affects soybean crops across the world (Wang et al. 2001). Multiple effector proteins have been reported for a plethora of plant pathogenic fungal and oomycete species (Selin et al. 2016).

The discovery of secreted effector proteins in these organisms was a major breakthrough, as these proteins are directly involved in pathogenicity. These effector proteins can be recognized by specific resistance genes (R genes) coding for proteins (Jones and Dangl 2006). The recognition of an effector protein by an R gene leads to effector triggered immunity (ETI), which generates a signaling cascade that results in programmed cell death via hypersensitive response (HR). This programed cell death slows the growth of the pathogen and avoids the proliferation of disease into neighboring plant cells (Jones and Dangl 2006).

Plant pathogenic oomycetes, or water molds, are a group of highly devastating plant pathogens. The diseases caused by these organisms can affect hundreds of plant species, and have led to high mortality of trees in forest ecosystems, losses of millions of dollars to agriculture, and been implicated in the Irish potato famine (Grünwald et al. 2008; Fry 2008). Widely recognized plant pathogen genera found in the water molds include *Phytophthora*, *Pythium*, *Albugo*, and *Peronospora* (Erwin and Ribeiro 1996).

Oomycetes contain a high number of predicted effector proteins (Birch et al. 2006; Kamoun 2006; Kamoun 2007). These proteins are secreted by the pathogen haustorium and translocated into the host plant cell, where they are transported to different organelles to disrupt physiological functions and facilitate disease (Birch et al. 2006; Kamoun 2006). Recent advances in molecular and computational biology have provided new information about the amino acid sequence of these oomycete effectors. Conserved motifs were identified in two key oomycete effector protein families: The RxLR-dEER motif for RxLR effectors, and the LFLAK-HVLV motif for crinkler (CRN) effectors. These canonical motifs are located near the N-terminus of an amino acid sequence following the signal peptide sequence. To date, hundreds of these RxLR and CRN effectors per species have been predicted for important plant pathogenic oomycetes from the genus *Phytophthora*, such as *Phytophthora infestans* (∼ 580 RxLR and ∼ 196 CRN effector genes (Haas et al. 2009)), *P. sojae* (∼ 470 RxLR and ∼100 CRN effector genes (Tyler et al. 2006)) and *P. ramorum* (∼ 260 RxLR and ∼19 CRN effector genes (Tyler et al. 2006)). In addition, the downy mildew pathogen *Hyaloperonospora arabidopsidis* contains up to 134 predicted RxLR-coding genes in its genome (Pel et al. 2014). However, not all oomycete species contain evidence of RxLR-coding genes in their genomes. No RxLR effector proteins have to date been found in the genera *Pythium, Saprolegnia,* or *Albugo* (Lévesque et al. 2010, Win et al. 2012, Links et al. 2011).

The process of identifying candidate effector-coding genes in sequenced genomes has, to date, been *ad ho*c and not fully reproducible. The most common bioinformatic process used for predicting effectors was described by Hass et al. (2009), where the authors used a combination of pattern matching via regular expressions (REGEX) based on the canonical RxLR and LFLAK motifs, followed by homology searches using Markov models. This method was used for the prediction of effector proteins in *Phytophthora infestans*, resulting in a total of 562 predicted RxLR effectors and 196 CRN effectors (Hass et al. 2009). The approach used by Hass et al. (2009) has been modified to include predicted effector sequences from other species, in order to improve the homology search (Govers and Bouwmeester 2008). Several independent tools are available to perform different steps to recognize effectors (Sonah et al. 2016, Dalio et al. 2017), but to date no bioinformatic tool allows the prediction of oomycete effectors in a simple and fast manner.

We developed the expandable *effectR* R package designed to predict effector proteins in a fast and reproducible way. This package uses both REGEX and homology search mechanisms based on Hidden Markov Models (HMM) to predict candidate effector proteins using gene models obtained from whole genome sequences. *effectR* provides functions in the statistical and computer language R (R Core Team 2016), which allow rapid identification and evaluation of candidate effector proteins for any oomycete species with a sequenced genome. We tested the *effectR* package using available oomycete genomes, and successfully validated effector prediction for the species *Phytophthora infestans*. The *effectR* package is modular and can be expanded to predict effector proteins with different canonical motifs specified by the user. In fact, we encourage contributions to our github repository of new functionality for effector prediction for other organismal groups. We included an example of a custom implementation for bacterial proteins that contain PAAR repeats. The *effectR* package is released on CRAN and a brief user tutorial is provided on github.

## RESULTS AND DISCUSSION

We developed the R package *effectR* that provides a convenient tool for rapid prediction of effectors in oomycete genomes, or with custom scripts for any genome, in a reproducible way. The *effectR* package relies on a combination of regular expressions statements and hidden Markov model approaches to predict candidate RxLR and CRN effectors using several function calls (Table 1). Three steps are required to call effectors: 1) A regular expression (REGEX) search to select amino acid sequences translated from all the open reading frames (ORFs) in a genome that match the motifs of interest, 2) a second, broad search of the amino acid sequences that match a Markov chain profile created with the amino acids that matched the motifs of interest in the REGEX step, and 3) a post-hoc set of tools combining the results from the two previous steps after filtering for redundancies (Figure 1). We are using the oomycete *P. infestans* as an example application:

**Table 1.** Functions found in *effectR* and their descriptions.

| Function | Description |
| --- | --- |
| `effector.summary()` | Returns non-redundant sequences from `hmm.search()` or `regex.search()` and generates a motif table. |
| `hmm.logo()` | Plots the relative frequencies of each position for the `hmmsearch()` table. |
| `hmm.search()` | Searching for motifs using HMM searches. |
| `regex.search()` | Searching for motifs using regular expressions (REGEX). |
| `shiny.effectR()` | A function to run *effectR* functions in a GUI using the shiny app. |

**Figure 1.**
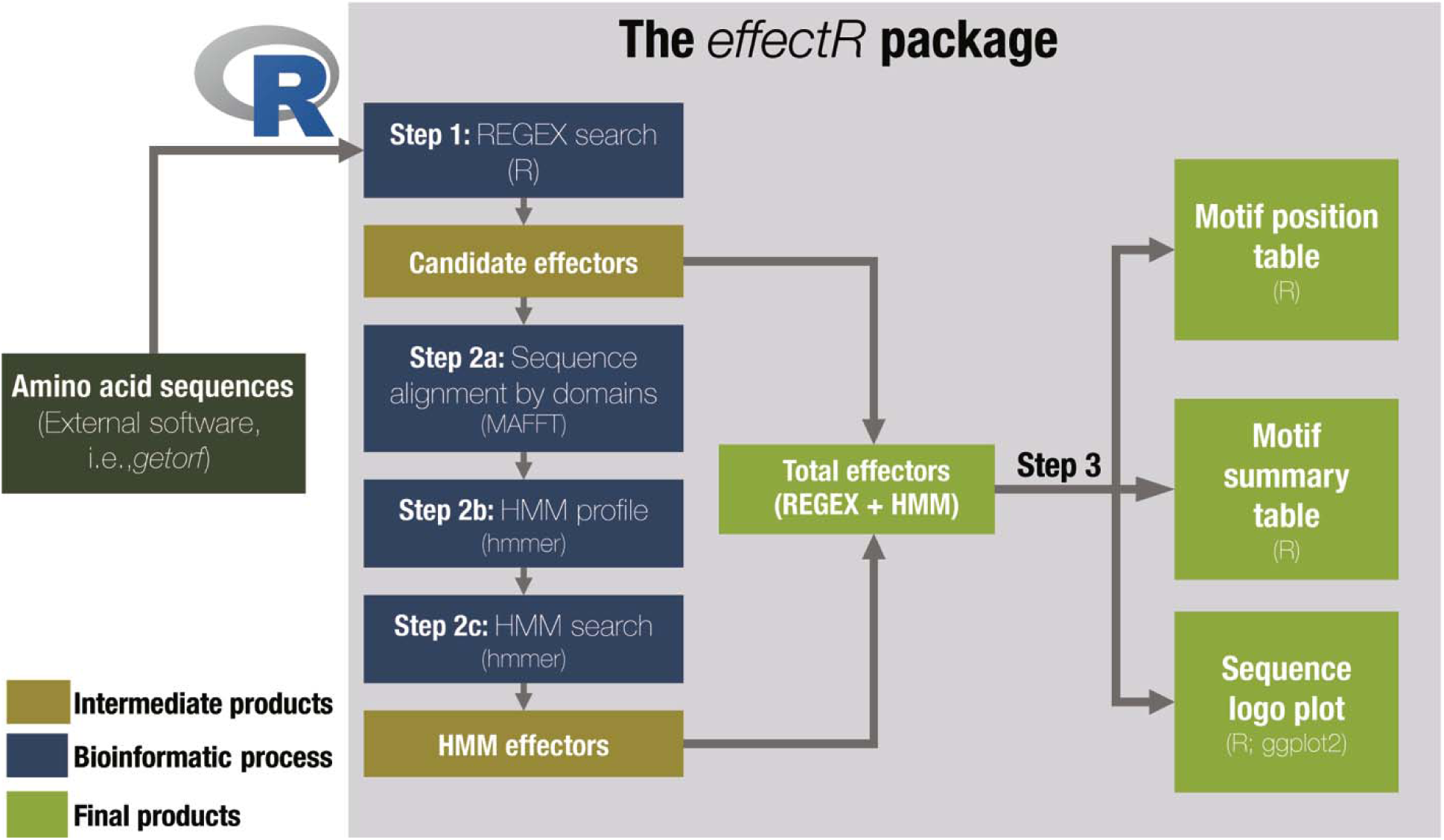
Workflow representing the steps used in the *effectR* R package. The package uses an amino acid FASTA file as input that contains either all translated gene models from a sequenced genome, or the 6-frame translation open reading frames (ORF) of a genome to obtain effectors. In this figure, the genomic ORFs represent 6-frame translations of a genome using the external software getorf from EMBOSS (Rice et al. 2000). After reading the amino acid FASTA input file into R, the *effectR* package predicts all effector proteins using three steps: 1) Searching MOTIF patterns via regular expression (REGEX), 2) searching for additional effectors using a homology search based on hidden Markov models using *hmmer*, and 3) a post-hoc analysis of the candidate effectors to visualize (Figure 4) and summarize the motifs of the candidates obtained by the package. All steps are modular and can be modified for discovery of any kind of effector.

### Step 1: Obtaining candidate RxLR and CRN effectors using regular expressions

The following example uses the function `regex.search()` with the test dataset test_infestans.fasta (a subset of 27 translated ORF’s of *P. infestans* from Haas et al. (2009) with 19 sequences with RxLR-EER motifs) included in the *effectR* package:

~~~
# Loading the effectR package in R
library(“effectR”)

# Using the read.fasta function of the seqinr package to import
the translated ORF FASTA file
fasta.file <-system.file("extdata", "test_infestans.fasta",
      package = "effectR")
ORF <-seqinr::read.fasta(fasta.file)
# Verifiyng if the length of the loaded FASTA file correspond to
the 27 translated ORF’s from P. infestans
length(ORF)

## [1] 27

# Executing step 1: Prediction of RxLR effectors from the ORF
object. Results are saved in the rxlr.cand object
rxlr.cand <-regex.search(ORF, motif = "RxLR")

# What is the number of RxLR effectors predicted by step 1?
length(rxlr.cand)

## [1] 18
~~~

This code snippet illustrates that `regex.search()` was able to predict 18 RxLR candidates out of the 27 translated ORF’s from the included test_infestans.fasta dataset. In addition, the *effectR* package can also predict crinkler effectors (CRN) or predict other families of interest based on custom regular expressions provided by the end user. Changing the `motif="RxLR"` option to `“CRN”` or `“custom”` will allow the prediction of these other motifs of interest without reloading the ORF dataset and significantly reducing processing time. Users are also encouraged to submit new functions to the *effectR* github repository for prediction of other motifs for any organismal group (see below and supplementary text 1 for an example).

### Step 2: Using Hidden Markov Models to predict additional candidate effectors

The following code example uses the function `hmm.search()` on the test_infestans.fasta dataset after obtaining the REGEX RxLR candidates:

~~~
# Loading the effectR package in R
library(“effectR”)

# Using the read.fasta function of the seqinr package to import
the translated ORF FASTA file
fasta.file <-system.file("extdata", "test_infestans.fasta",
        package = "effectR")
ORF <-seqinr::read.fasta(fasta.file)

# Step 1 prediction
REGEX <-regex.search(ORF, motif = "RxLR")

# Expanding the search of RxLR effectors using HMM searches (step
2). All candidate effectors predicted by step 2 will be saved in
the candidate.rxlr object
candidate.rxlr <-hmm.search(original.seq = fasta.file, regex.seq
      = REGEX)

## Starting MAFFT alignment.
## ---
## Executing MAFFT
## Please be patient
## MAFFT alignment finished!
## Starting HMM ## ---
## Creating HMM profile

[additional console output was removed for brevity]

## HMM search done.
## ---
##
## Total of sequences found in REGEX: 18
## Total of sequences found in HMM: 19
## Total of redundant hits: 18
## Number of effector candidates: 19
~~~

The previous code snippet shows that `hmm.search()` was able to identify one new RxLR candidate from the test_infestans.fasta dataset, increasing our total number of candidate RxLR effectors to 19. The `hmm.search()` function also detects candidate effectors previously predicted by the REGEX step. In the previous example, the `hmm.search()` function detected 18 candidate effectors using the `regex.search()` step. The combined 19 candidate effectors based on the REGEX and HMM searches can now be evaluated in step 3.

### Step 3: Post-hoc tools for curation of candidate effector genes

In the following steps, *effectR* provides functions to summarize and visualize candidate effector genes.

#### Step 3a: Summarizing the predicted effector proteins and determining the number and position of each motif in the effector sequences

The *effectR* package includes the `effector.summary()` function that combines the results from both REGEX and HMM searches into a list of unique predicted effectors that can be exported into multiple sequence format files. The `effector.summary()` function returns a table of the number of motifs per sequence and the position of the first residue of each motif of interest for all candidate effectors. In addition, the table includes a “summary motif” column that will define the candidate sequence as: “Complete” (includes both motifs of interest), “Only × motif” (if the sequence includes only one of the two motifs of interest), or “No Motifs” (No motifs of interest found):

~~~
# Loading the effectR package in R
library(“effectR”)
# Using the read.fasta function of the seqinr package to import
the translated ORF FASTA file
fasta.file <-system.file("extdata", "test_infestans.fasta",
         package = "effectR")
ORF <-seqinr::read.fasta(fasta.file)

# Step 1 prediction
REGEX <-regex.search(ORF, motif = "RxLR")

# Expanding the search of RxLR effectors using HMM searches (step
2). All candidate effectors predicted by step 2 will be saved in
the candidate.rxlr object
candidate.rxlr <-hmm.search(original.seq = fasta.file, regex.seq
       = REGEX)

# Summarizing the predictions from step 1 and step 2 using the
effectR.summary() function (Step 3a).

# The summary of non-redundant RxLR predicted effectors will be
stored in the RxLR.effector$consensus.sequences objects

#The table of motif number and position will be stored in the
RxLR.effectors$motif.table object
RxLR.effectors <-effector.summary(candidate.rxlr, motif =
       "RxLR")

# What is the number of non-reduntant RxLR effectors predicted by step
          1 and step 2?
length(RxLR.effectors$consensus.sequences)

## [1] 18

# Table of the first five motif number and position for all non-
redudant, predicted RxLR effector proteins
head(RxLR.effectors$motif.table, n = 5)

##Sequence ID                 RxLR number  RxLR position  EER number     EER position   MOTIF
##DS028118_1098 DS028118_1098     1              44           1             54         Complete
##DS028118_4814 DS028118_4814     1              40           1             53         Complete
##DS028118_5398 DS028118_5398     1              39           1             49         Complete
##DS028120_2026 DS028120_2026     1              42           1             53         Complete
##DS028120_2220 DS028120_2220     1              41           1             55         Complete
~~~

#### Step 3b: Plotting the HMM profile

To visualize the position of the motifs of interest, we include the function `hmm.logo()` that plots the results from the HMM profile obtained in *hmmbuild* in a logo plot (Schneider and Stephens 1990) using the *ggplot2* package (Wickham 2016). The `hmm.logo()` function reads the *hmmer* profile table and extracts a bit score of each amino acid at each position. Then, *effectR* plots the bit score on the Y axis, the amino acid position in the × axis, and overlays the amino acid with the highest bit score over the plot in order to represent the frequency for each amino acid found at every position of the consensus sequence from the multiple sequence alignment step. Here’s an example of the `hmm.logo()` function for the *P. infestans* example dataset:

~~~
# Loading the effectR package in R
library(“effectR”)

# Using the read.fasta function of the seqinr package to import
the translated ORF FASTA file
fasta.file <-system.file("extdata", "test_infestans.fasta",
      package = "effectR")
ORF <-seqinr::read.fasta(fasta.file)

# Step 1 prediction
REGEX <-regex.search(ORF, motif = "RxLR")

# Expanding the search of RxLR effectors using HMM searches (step
2). All candidate effectors predicted by step 2 will be saved in
the candidate.rxlr object
candidate.rxlr <-hmm.search(original.seq = fasta.file, regex.seq
      = REGEX)

# Plotting the HMM profile created in the hmm.search() function
using the hmm.logo() function
hmm.logo(candidate.rxlr$HMM_Table)

## R graphical output [NOTE: NOT SHOWN HERE, BUT PRINTED TO
CONSOLE]:
~~~

### Validation

To test the developed package, we used a set of four sequenced genomes of three oomycete species and one fungal species to predict RxLR effectors. We used the genomes of the oomycete species *Phytophthora infestans* (Haas et al. 2009), *Pythium ultimum* (Lévesque et al. 2010), and *Albugo candida* (Links et al. 2011). *P. infestans* has more than 580 predicted RxLR effectors (Haas et al. 2009), while neither *P. ultimum* nor *A. candida* have any reported RxLR motifs. Instead, *A. candida* has 26 predicted gene models with a variant RxLR motif called Ac-RXL (Links et al. 2011). The genome of the ascomycete *Fusarium oxysporum* f. sp. *lycopersici* (Ma et al. 2010) was used as a negative control or outgroup with no expectation of finding RxLR motives.

The assembled reference genomes of each of the 4 species were downloaded from Fungi-DB (Stajich et al. 2011). For each genome assembly, a six-frame translation of all ORFs was predicted using *getorf* from EMBOSS (Rice et al. 2000). Translated ORF’s with a length of less than 100 amino acid residues were discarded. We predicted all RxLR effector proteins for each translated sequence from the genome assembly in the *effectR* package. Additionally, prediction of the signal peptide using all predicted candidate effectors was performed in SignalP 3.0 (Bendtsen et al. 2004). A threshold D-score > 0.8 was used. Any candidate effector with a predicted signal peptide was considered a valid candidate effector. All valid candidate effector proteins from *P. infestans* were compared to the published list of effectors from Haas et al. (2009). We created a custom blast database using the RxLR effector proteins and identified matches between the amino acid sequences of the *effectR* predictions against the RxLR candidate database in *blastp* (Altschul et al. 1990).

A total of 174 candidate effectors were predicted for *P. ultimum*, 47 for *A. candida*, and 3 for *F. oxysporum* f. sp. *lycopersici*. None of these predicted effectors were considered valid, as no evidence of a signal peptide cleavage site was found for any of the predicted sequences (Table 2). These results are consistent with previous reports of finding no RxLR effector proteins in genomes of these oomycete and fungal species. The Ac-RXL effector proteins of *A. candida* appear to have an independent origin than the RxLR effector proteins found in *Phytophthora* species (Links et al. 2011), and no functional RxLR effector proteins have been reported for *Fusarium oxysporum* f. sp. *lycopersici* (Ellis et al. 2009; Ma et al. 2010).

**Table 2.** Predicted RxLR effector proteins in *effectR* for the open reading frame translations (ORF) of the assembled genomes of *Phytophthora infestans, Pythium ultimum, Albugo candida, and Fusarium oxysporum f. sp. lycopersici.* The table includes the total number of predicted effector proteins at each step of the package (step 1: REGEX search; step 2: HMM search) and the total number of unique effector proteins (intersection of step 1 with step 2). If the candidate effector protein obtained in *effectR* does not have a signal peptide, then the predicted RxLR effector protein is not considered valid.

| Species | Number of<br>predicted ORF | Candidate RxLR effector<br>proteins<br>predicted by *effectR* |  |  | Number of valid RxLR effectors<br>protein with a signal peptide<br>cleavage site |
| --- | --- | --- | --- | --- | --- |
|  |  | REGEX | HMM | Total |  |
| *Phytophthora infestans* | 208,272 | 395 | 827 | 900 | 631 |
| *Pythium ultimum* | 63,882 | 61 | 149 | 174 | 0 |
| *Albugo candida* | 24,405 | 13 | 34 | 47 | 0 |
| *Fusarium oxysporum* f. sp.<br>*lycopersici* | 55,224 | 3 | 0 | 3 | 0 |

For *P. infestans*, we predicted 395 candidate effector proteins in the REGEX step, and 827 candidate effector proteins in the HMM step, for a total of 900 non-redundant RxLR effector proteins (a number similar to the 831 RxLR effectors predicted by Haas et al. (2009; see Haas et al. Supplementary Methods 2) using only a combination of REGEX and HMM methods). In contrast to the predictions in the previously screened fungal and oomycete species, a high number of valid effectors (631 valid candidate effectors with evidence of a signal peptide cleavage site) were predicted for the translated sequences of *P. infestans* (Table 2). The logo plot shows the prevalence of the RxLR-EER motifs obtained by the REGEX step between residues 49 and 67 (Figure 4). The high number of effector proteins predicted in the HMM step is a result of the low thresholds used by our package in order to obtain as many candidate effectors as possible. Out of the 631 valid candidate effector proteins, we predicted 453 RxLR effectors with a *blastp* match of more than 90% identity with the Haas et al. (2009) RxLR predicted effector proteins. The genomic position of each of these 453 predicted proteins also corresponded with the position of the homologous effector protein reported by Haas et a. (2009), indicating that *effectR* successfully predicted previously known RxLR effector proteins from *P. infestans*. Our approach did not predict 99 predicted effector proteins from Haas et al. (2009). Out of these 99 RxLR effector proteins, we find 18 proteins with no presence of either an RxLR or EER domain, 24 proteins with no evidence of EER domain and the remainder 57 proteins with the RxLR/EER domain with higher upstream/downstream distances from the REGEX expected residue position. These results are expected as Haas et al. (2009) used a combination of *blastp* searches, TribeMCL clustering and homology searches from reference effectors in addition to the REGEX + HMM searches, leading to a higher number of candidate effector proteins than expected just from the searches implemented in this package. *effectR* also predicted 79 candidate effector proteins not present in Hass et al. (2009) that include evidence of a signal peptide cleavage site (Table 3). Out of these 79 newly predicted effector proteins, 40 proteins contained both RxLR and EER domains, 14 only contained the RxLR motif, 18 only had the EER motif, and 7 proteins did not include any of the motifs of interest (Table 3).

**Table 3.** Seventy-nine new candidate RxLR effector proteins from *P. infestans*  predicted by the *effectR* package. None of these candidate proteins were predicted by Haas et al. (2009). The candidate effector proteins are sorted by scaffold and start position. “Complete” refers to candidate RxLR effector proteins with both RxLR and EER domains present in the amino acid sequence.

| Scaffold | Start | End | Number of<br>RxLR<br>motifs | Position of<br>RxLR motif | Number of<br>EER motifs | Position of<br>EER motifs | Motifs present |
| --- | --- | --- | --- | --- | --- | --- | --- |
| Supercontig_1.2 | 387,181 | 387,789 | 0 | NA | 0 | NA | No motifs |
| Supercontig_1.2 | 5,134,349 | 5,134,744 | 1 | 35 | 1 | 51 | Complete |
| Supercontig_1.3 | 4,668,546 | 4,669,019 | 1 | 43 | 1 | 66 | Complete |
| Supercontig_1.4 | 1,947,482 | 1,947,706 | 0 | NA | 0 | NA | No motifs |
| Supercontig_1.5 | 2,020,297 | 2,020,617 | 1 | 40 | 0 | NA | Only RxLR motif |
| Supercontig_1.5 | 3,399,390 | 3,400,325 | 0 | NA | 1 | 47 | Only EER motif |
| Supercontig_1.5 | 1,365,830 | 1,366,261 | 1 | 54 | 1 | 81 | Complete |
| Supercontig_1.5 | 3,296,675 | 3,297,394 | 1 | 31 | 1 | 44 | Complete |
| Supercontig_1.5 | 1,695,450 | 1,695,806 | 1 | 43 | 1 | 56 | Complete |
| Supercontig_1.7 | 778,069 | 778,482 | 1 | 50 | 0 | NA | Only RxLR motif |
| Supercontig_1.8 | 2,694,681 | 2,694,974 | 0 | NA | 0 | NA | No motifs |
| Supercontig_1.9 | 417,751 | 418,602 | 1 | 47 | 0 | NA | Only RxLR motif |
| Supercontig_1.10 | 726,219 | 726,617 | 0 | NA | 1 | 59 | Only EER motif |
| Supercontig_1.11 | 1,535,262 | 1,536,773 | 0 | NA | 0 | NA | No motifs |
| Supercontig_1.11 | 1,187,840 | 1,188,091 | 1 | 50 | 1 | 63 | Complete |
| Supercontig_1.11 | 1,786,307 | 1,786,816 | 1 | 50 | 1 | 60 | Complete |
| Supercontig_1.11 | 529,720 | 529,944 | 1 | 41 | 0 | NA | Only RxLR motif |
| Supercontig_1.11 | 1,902,450 | 1,903,310 | 1 | 53 | 1 | 61 | Complete |
| Supercontig_1.13 | 612,373 | 612,726 | 1 | 50 | 1 | 63 | Complete |
| Supercontig_1.14 | 2,070,861 | 2,072,003 | 2 | 57,322 | 1 | 73 | Complete |
| Supercontig_1.15 | 1,593,048 | 1,593,419 | 0 | NA | 1 | 62 | Only EER motif |
| Supercontig_1.15 | 668,809 | 669,123 | 1 | 23 | 1 | 45 | Complete |
| Supercontig_1.15 | 2,682,075 | 2,682,359 | 1 | 49 | 0 | NA | Only RxLR motif |
| Supercontig_1.16 | 1,715,898 | 1,716,353 | 1 | 65 | 0 | NA | Only RxLR motif |
| Supercontig_1.16 | 1,567,608 | 1,568,108 | 1 | 57 | 1 | 76 | Complete |
| Supercontig_1.17 | 448,981 | 449,679 | 1 | 47 | 1 | 83 | Complete |
| Supercontig_1.18 | 806,113 | 806,700 | 1 | 43 | 1 | 51 | Complete |
| Supercontig_1.20 | 1,412,054 | 1,413,028 | 0 | NA | 1 | 62 | Only EER motif |
| Supercontig_1.20 | 1,339,618 | 1,340,277 | 1 | 52 | 0 | NA | Only RxLR motif |
| Supercontig_1.25 | 1,219,557 | 1,220,138 | 0 | NA | 1 | 60 | Only EER motif |
| Supercontig_1.25 | 1,023,691 | 1,023,990 | 0 | NA | 1 | 46 | Only EER motif |
| Supercontig_1.25 | 1,314,779 | 1,315,240 | 1 | 42 | 1 | 56 | Complete |
| Supercontig_1.25 | 1,633,314 | 1,633,970 | 1 | 48 | 1 | 68 | Complete |
| Supercontig_1.26 | 1,717,796 | 1,718,344 | 0 | NA | 0 | NA | No motifs |
| Supercontig_1.27 | 816,842 | 817,168 | 1 | 50 | 1 | 67 | Complete |
| Supercontig_1.27 | 638,168 | 638,521 | 1 | 51 | 1 | 65 | Complete |
| Supercontig_1.29 | 807,454 | 807,804 | 0 | NA | 1 | 56 | Only EER motif |
| Supercontig_1.29 | 1,420,380 | 1,420,796 | 1 | 42 | 1 | 57 | Complete |
| Supercontig_1.32 | 40,829 | 41,050 | 1 | 47 | 1 | 64 | Complete |
| Supercontig_1.33 | 857,150 | 858,472 | 1 | 40 | 1 | 57 | Complete |
| Supercontig_1.33 | 1,310,346 | 1,310,840 | 1 | 56 | 0 | NA | Only RxLR motif |
| Supercontig_1.33 | 1,324,452 | 1,324,946 | 1 | 40 | 1 | 57 | Complete |
| Supercontig_1.34 | 80,145 | 80,615 | 0 | NA | 1 | 64 | Only EER motif |
| Supercontig_1.34 | 708,534 | 709,043 | 1 | 47 | 1 | 66 | Complete |
| Supercontig_1.35 | 875,543 | 876,028 | 1 | 46 | 0 | NA | Only RxLR motif |
| Supercontig_1.38 | 848,659 | 849,336 | 0 | NA | 0 | NA | No motifs |
| Supercontig_1.38 | 794,856 | 797,288 | 1 | 216 | 1 | 46 | Complete |
| Supercontig_1.39 | 1,376,318 | 1,376,599 | 1 | 50 | 1 | 63 | Complete |
| Supercontig_1.40 | 730,075 | 730,683 | 0 | NA | 1 | 62 | Only EER motif |
| Supercontig_1.42 | 695,078 | 695,443 | 1 | 30 | 0 | NA | Only RxLR motif |
| Supercontig_1.43 | 780,095 | 780,469 | 1 | 50 | 1 | 69 | Complete |
| Supercontig_1.43 | 1,272,359 | 1,272,643 | 1 | 53 | 1 | 63 | Complete |
| Supercontig_1.43 | 1,286,094 | 1,286,504 | 1 | 53 | 1 | 69 | Complete |
| Supercontig_1.44 | 865,765 | 866,316 | 0 | NA | 1 | 76 | Only EER motif |
| Supercontig_1.44 | 1,006,809 | 1,007,273 | 1 | 39 | 1 | 56 | Complete |
| Supercontig_1.47 | 1,060,823 | 1,061,302 | 1 | 54 | 1 | 74 | Complete |
| Supercontig_1.63 | 438,688 | 438,969 | 0 | NA | 1 | 69 | Only EER motif |
| Supercontig_1.63 | 549,263 | 549,847 | 0 | NA | 1 | 68 | Only EER motif |
| Supercontig_1.66 | 372,213 | 372,752 | 0 | NA | 1 | 76 | Only EER motif |
| Supercontig_1.66 | 259,626 | 260,069 | 1 | 44 | 1 | 56 | Complete |
| Supercontig_1.67 | 516,713 | 517,306 | 0 | NA | 1 | 64 | Only EER motif |
| Supercontig_1.79 | 318,551 | 319,021 | 1 | 51 | 0 | NA | Only RxLR motif |
| Supercontig_1.89 | 375,480 | 376,355 | 1 | 35 | 0 | NA | Only RxLR motif |
| Supercontig_1.89 | 370,258 | 370,542 | 1 | 52 | 1 | 60 | Complete |
| Supercontig_1.124 | 94,888 | 95,385 | 1 | 47 | 1 | 60 | Complete |
| Supercontig_1.129 | 11,438 | 16,690 | 1 | 53 | 1 | 67 | Complete |
| Supercontig_1.130 | 54,647 | 54,997 | 0 | NA | 1 | 56 | Only EER motif |
| Supercontig_1.141 | 177,814 | 179,079 | 0 | NA | 1 | 52 | Only EER motif |
| Supercontig_1.220 | 51,445 | 51,846 | 0 | NA | 1 | 61 | Only EER motif |
| Supercontig_1.231 | 92,953 | 93,969 | 1 | 38 | 1 | 49 | Complete |
| Supercontig_1.235 | 17,662 | 20,670 | 1 | 52 | 1 | 71 | Complete |
| Supercontig_1.241 | 140,057 | 140,608 | 0 | NA | 1 | 60 | Only EER motif |
| Supercontig_1.241 | 90,165 | 90,533 | 1 | 53 | 1 | 67 | Complete |
| Supercontig_1.302 | 59,522 | 59,977 | 2 | 51,122 | 1 | 70 | Complete |
| Supercontig_1.385 | 9,167 | 10,102 | 1 | 216 | 1 | 46 | Complete |
| Supercontig_1.709 | 2,672 | 2,869 | 0 | NA | 0 | NA | No motifs |
| Supercontig_1.1192 | 4,030 | 5,904 | 3 | 51,77,310 | 0 | NA | Only RxLR motif |
| Supercontig_1.2669 | 2,032 | 2,460 | 1 | 27 | 1 | 43 | Complete |
| Supercontig_1.3033 | 1,747 | 2,109 | 2 | 74,80 | 0 | NA | Only RxLR motif |

**Figure 4.**
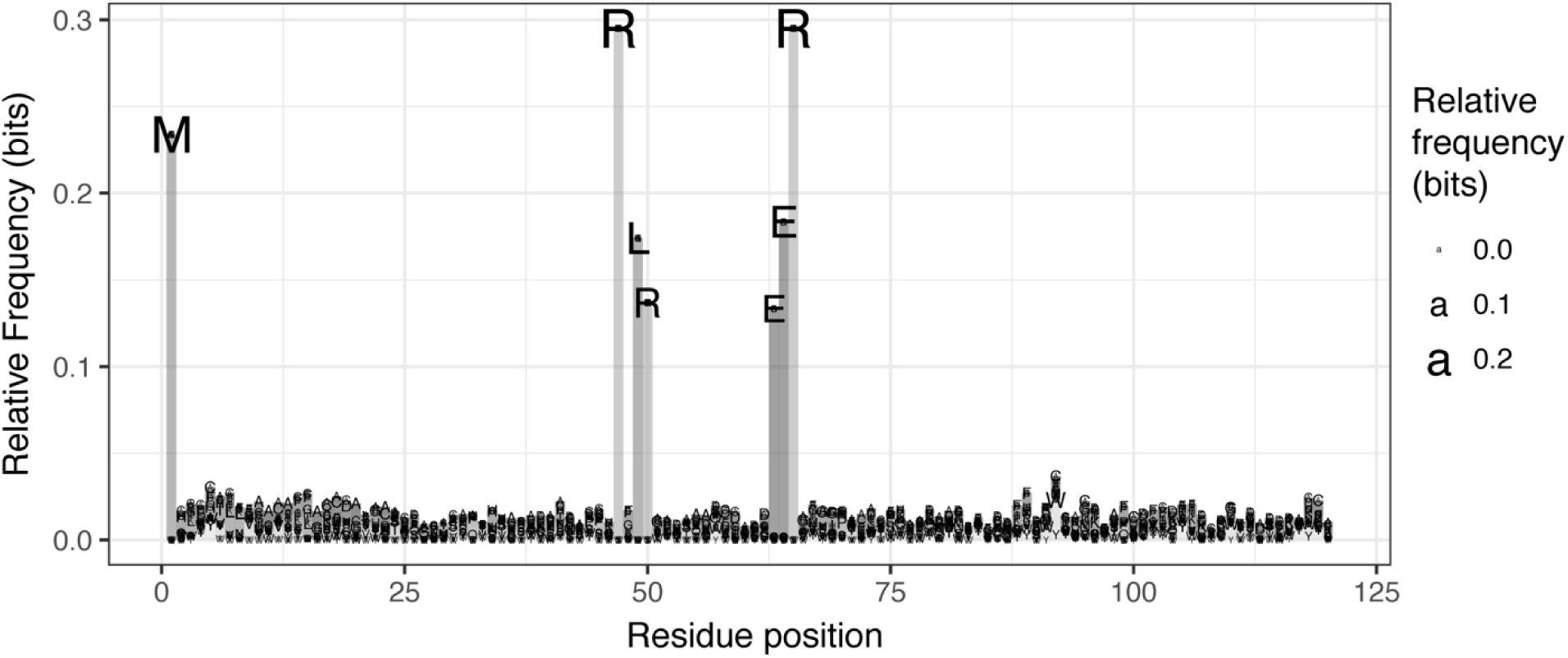
Sequence logo plot of the candidate RxLR’s obtained for *Phytophthora infestans* in the HMM profile built from the sequences obtained in the REGEX step (step 1). The size of the letters is proportional to the height of the bar, and it reflects the relative frequency of the particular amino acid. The plot also shows the high relative frequency of the RxLR-EER motifs around the 50th residue of the amino acid sequence.

In addition to the prediction of RxLR effector proteins, we predicted the CRN effector proteins for the ORF translations of the species used in the proof of concept (Supplementary Table 1). The results of the CRN prediction are consistent with the expectation of a high number of CRN effector proteins present in the genomes of oomycete species, with 21 candidate CRN effectors predicted for *P. ultimum* and 214 CRN effectors predicted for *P. infestans*. The number of predicted effectors for these two species are similar to the reported number of CRN effectors (196 CRN effectors for *P. infestans* and 26 CRN effectors for *P. ultimum*) (Lévesque et al. 2010). However, only four CRN effectors in *A. candida* were predicted. These four predicted CRN proteins for *A. candida* only have the LxLAK motif, as reported by Links et al. (2011) (Supplementary Table 1) and are not considered canonical CRN effectors (Links et al. 2011). Finally, only three CRN effectors were predicted for the fungal species *Fusarium oxysporum* f. sp. *lycopersici.* No CRN effectors have reported for this fungal pathogen, and the CRN effectors predicted by *effectR* only contain the LxLAK motif, indicating that these proteins are not canonical CRN effectors as in *A. candida.*

### Custom scripts for other effectors

*effectR* can easily be modified for identifying candidate effectors. For example, `regex.search()` can be modified to call bacterial proteins that contain PAAR repeats. These proteins are associated with the VgrG-like spikes found in the type 6 secretion system (T6SS) of bacteria and have been shown to be essential in target cell killing by the bacterial species *Vibrio cholerae* and *Acitenobacter baylyi* (Shneider et al. 2013). The PAAR proteins have a homonymous amino acid sequence motif (PAAR) with one or more repeats. We created an example how the *effectR* can predict potential candidate proteins in the proteome of the reference strain ATCC 39315 of *V. cholerae* (Heidelberg et al. 2000) by using the PAAR motif as part of the REGEX search (Supplementary text 1). Our results indicate that *effectR* successfully predicted 19 candidate PAAR proteins, two of which are homologous to the previously reported PAAR proteins. These two predicted proteins only differ in one amino acid when compared to PAAR homolog proteins previously described by Shneider et al. (2013) from other strains of *V. cholera*e (Supplementary text 1, Supplementary figure 1). These results show that *effectR* can correctly predict different proteins with other motifs than the canonical oomycete effectors, and also illustrate the importance of using manual curation based on homologous proteins to avoid the detection of false positives.

### Conclusions

Our *effectR* package provides a novel tool for reproducible prediction of candidate effectors in oomycete genomes. The package is modular, where every step can be modified to predict candidate effector proteins. Custom motifs for any new effector family can easily be added via custom scripts or by contributing to the github repository. The package has been successfully tested for translated ORFs predicted from genomes of oomycete plant pathogens and can be used for a quick survey of the effector arsenal involved in any plant-pathogen interactions for any species of interest where effector motifs are known.

## MATERIALS AND METHODS

### The *effectR* package

The *effectR* package is written in the R computer language (R Core Team 2018). *effectR* allows prediction of oomycete effectors using three steps (Figure 1): 1) A regular expression (REGEX) search to select amino acid sequences translated from all the open reading frames (ORFs) in a genome that match the motifs of interest, 2) a second, broad search of the amino acid sequences that match a Markov chain profile created with the amino acids that matched the motifs of interest in the REGEX step, and 3) a *post-hoc* set of tools combining the results from the two previous steps after filtering for redundancies. The package requires as input a FASTA file ideally containing all six-frame amino acid translations for each open reading frame of the sequenced genome of interest or at a minimum all translated gene models. The package then returns the total number of predicted effectors, the amino acid sequence for each of the predicted effectors, the number and position of the motifs of interest for each predicted effector, and the Markov chain profile table which can be conveniently visualized.

### External programs required to execute *effect*R

The *effectR* package requires installation of additional programs. *effectR* can use amino acid translations of the gene models predicted in a genome of interest, but we recommend the use of six-frame translations of all open reading frames (ORF) from a genome assembly. Including all six translations for each ORF in a genome will allow prediction of more candidate effectors present in an organism of interest. To generate six-frame ORF translations we use *getorf* from EMBOSS (Rice et al. 2000). *getorf* can be run locally on the command line or online via the EMBOSS explorer (Rice et al. 2000). Other additional programs required by *effectR* are the MAFFT v7 multiple sequence alignment tool (Katoh and Standley 2013) and HMMER 3.1b2 (Eddy 2011). MAFFT performs a sequence alignment of the candidate effectors in order to build a reference profile to be scanned by HMMER. HMMER executes the searches based on hidden Markov models (Figure 1, step 2). These two programs are external to R and must be installed on the same machine as *effectR*. The *effectR* package includes functions to detect if both HMMER and MAFFT are available in the default user path or the user can specify the location of each of the binaries when executing *effectR*.

While these additional programs are indispensable to complete step 2 of the *effectR* package, step 1 and step 3 can be executed independently of them. The user can execute step 1 within *effectR*, use the output from step 1 and the original ORF file to perform a multiple sequence alignment (MSA) using other tools of interest, and execute HMM searches externally. The *effectR* package can import external MSA in FASTA format and HMM results in table format to be used in step 3. The inclusion of step 2 as part of the package was created for a more streamlined process in which the user can identify candidates using REGEX searches, broaden the number of candidate proteins via HMM searches, and summarize and quality control the results obtained from the *effectR* package in a fast and seamless manner.

### Obtaining candidate RxLR and CRN effectors using regular expressions

To predict the first set of candidate effector proteins, *effectR* searches the ORF translation file to find sequences that match the motifs of interest (Figure 1). These searches are based on regular expression matching. For the RxLR motif, the regular expression (REGEX) reported by Haas et al. (2009) is used:

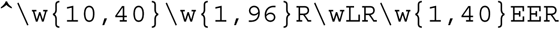

This REGEX example shows the three parts used in step 1 to identify an RxLR effector candidate (Figure 2): Part 1A (\w{10,40}) reserves the first 10 to 40 positions of the amino acid sequence for the signal peptide. Part 1B (\w{1,96}R\wLR) searches for amino acid residues that matches the RxLR motif within the following 1-96 residues after the signal peptide. Part 1C (\w{1,40}EER) searches for the EER motif within the 40 residues following the RxLR motif. A limitation of *effectR* is that it does not directly predict the presence of a signal peptide. Other programs (i.e., SignalP 3.0 (Bendtsen et al. 2004)) can be used to predict the presence of the signal peptide.

**Figure 2.**
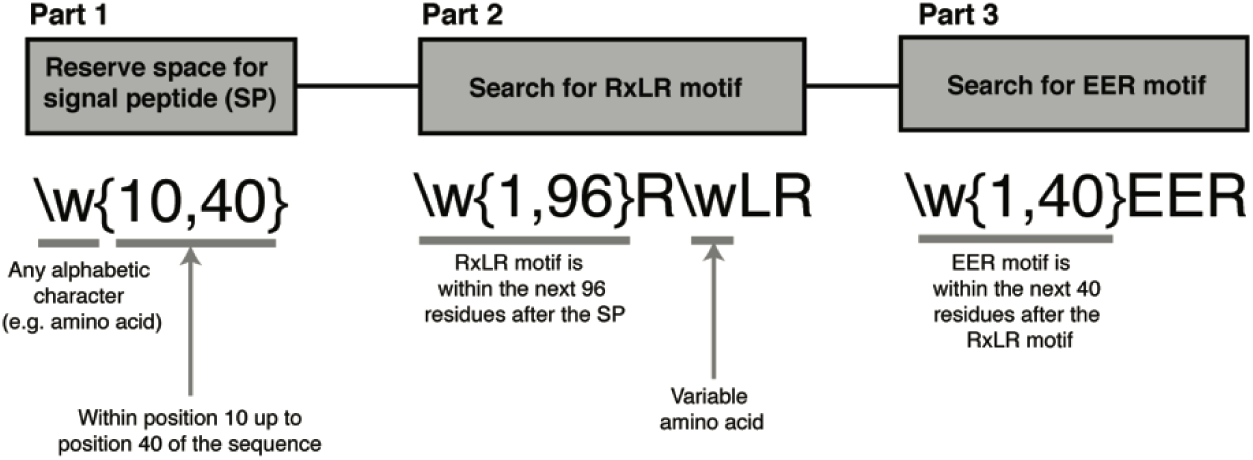
Graphical representation of the regular expression (REGEX) used by *effectR* to predict candidate RxLR effectors. This REGEX is integrated in the `regex.search()` function of *effectR*. The REGEX search is divided into 3 parts (grey boxes). Part 1: reserves the space for the signal peptide; part 2: performs a downstream search for the RxLR motif; and part 3: searches for the EER motif downstream from the RxLR motif. If a query amino acid sequence has a match to the two motifs, the sequence is considered a candidate RxLR effector protein, and used to build the HMM profile for step 2 of the package.

Step 1 is summarized in the `regex.search()` function of the *effectR* package. Step 1 determines if a translated ORF is a candidate effector after identifying the presence of a RxLR/CRN motif via REGEX (RxLR motif patterns:

`^\w{10,40}\w{1,96}R\wLR\w{1,40}EER, CRN` motif pattern:

`^\w{1,90}LFLAK\w+` (Haas et al. 2009)). In addition, we have added a “custom” option that allows user-specification of a custom regular expression, in order to identify candidate genes for other effectors of interest.

### Using Hidden Markov Models (HMM) to predict additional candidate effectors

A second independent method of identifying candidate effectors relies on HMM (Figure 1, step 2). This search allows the identification of additional sequences that match a HMM profile. In *effectR*, the HMM profile is built using the candidate effectors predicted in the REGEX step. The HMM profile includes the probabilities of any amino acid occurring in a given position of the consensus sequence. The consensus sequence is the product of a multiple sequence alignment of the sequences used to build the HMM profile (Eddy 1998).

To create the HMM profile, *effectR* aligns the candidate effectors to identify common motifs and builds the HMM profile based on these common motifs. *effectR* uses MAFFT (Katoh and Standley 2013) to calculate the multiple sequence alignment of the candidate effectors predicted from the REGEX step. The package *effectR* uses the E-INS-i iterative refinement algorithm for creating the alignment. The E-INS-i algorithm is suitable for sequences containing common domains of interest flanked by large un-alignable regions (Katoh and Standley 2013). These large, un-alignable regions are typically observed in RxLR and CRN motifs. After generating the multiple sequence alignment from REGEX candidate effectors, *effectR* creates a HMM profile using HMMER’s *hmmbuild* and *hmmpress* modules. After the HMM profile is built, *effectR* searches the original ORF file for sequences with hits against the HMM profile using hmmscan. To encourage the user to perform manual curation, *effectR* does not apply any significance thresholds in the *hmmsearch* step and returns all of the sequences that match the HMM profile to the user. The result from the HMM search is a list of translated ORFs that match the HMM profile. This list can be used for manual curation steps (Figure 1, step 3) or can be exported using the write.fasta function of the *seqinr* package (Charif and Lobry 2007). This step has been summarized in the `hmm.search()` function of the *effectR* package.

### Post-hoc tools for combining candidate effector genes from REGEX and HMM searches

The *effectR* package allows for manual curation of the predicted effector proteins to screen for highly heterogeneous sequences or for sequences that match the HMM profile but do not include any of the motifs of interest in their amino acid sequence. The manual curation functions created for the *effectR* package includes the `effector.summary()` function. This function summarizes the results from step 1 and step 2 and generates a list of unique candidate effectors. The `effector.summary()` function also returns the numbers of motifs and position in the amino acid sequence for all predicted candidate effectors. Finally, we created the `hmm.logo()` function to plot logo-like figures based on the candidate effectors predicted in steps 1 and 2. Each step is modular and fully compatible with the results of either REGEX or HMM searching.

### Software availability

The *effectR* package can be downloaded from CRAN or GitHub as described at https://github.com/grunwaldlab/effectR. *effectR* can be used on the command line or can be deployed in a user-friendly, point-and-click graphical interface via the shiny R framework (Chang et al. 2017) using the shiny.effectR() function (Figure 3). The end user has to install the packages *shiny* (Chang et al. 2017) and *shinyjs* (Attali 2018) in order to deploy the shiny R graphic interface.

**Figure 3.**
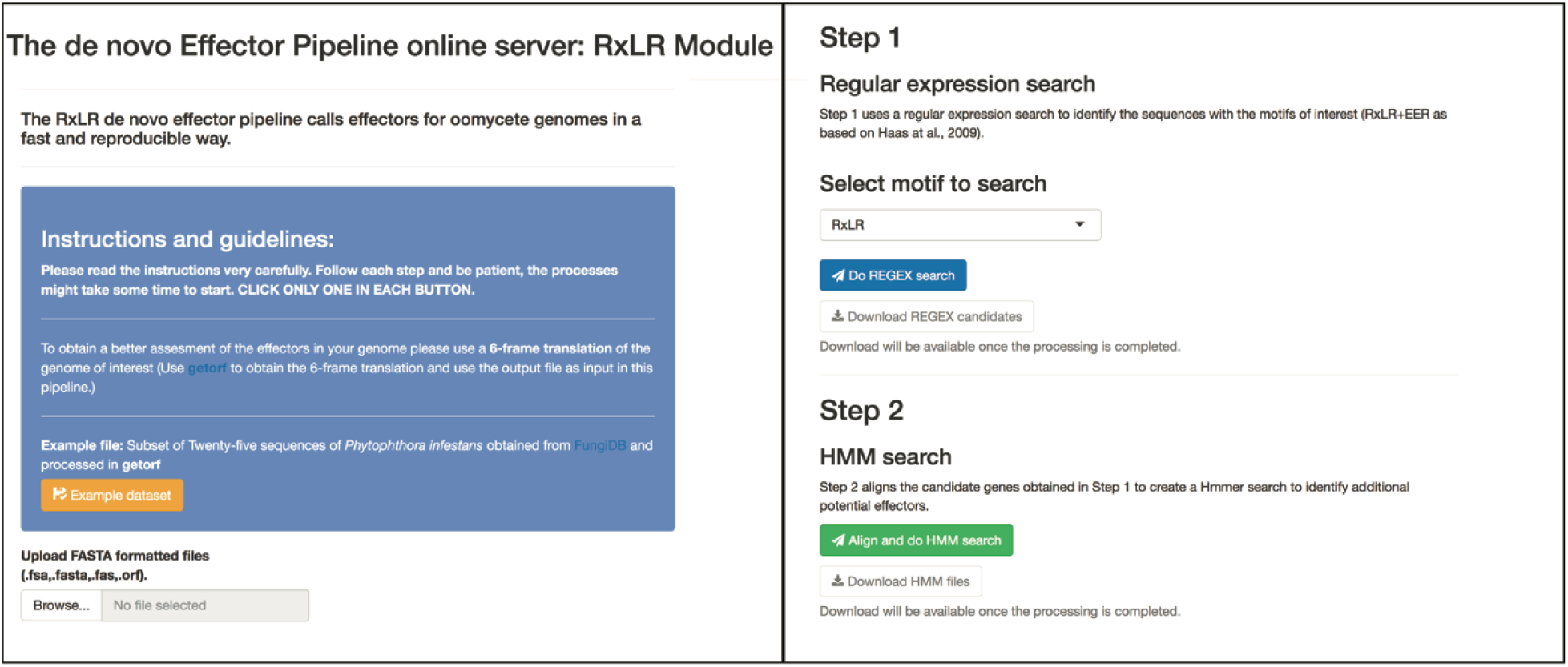
Screen capture of the Shiny R web-application for *effectR*. All functions available in the package are included in this web-application. The web-application can be initiated within an active R session by running the `shiny.effectR()` function.

## ACKNOWLEDGMENTS

We thank Brian Knaus, Zhian Kamvar, and Zach Foster for their technical advice, extensive code review, and vast expertise in R programing. This work was supported by funds from USDA ARS Project 2072-22000-041-00-D and USDA-NIFA Project 2010-511001-21649. Mention of trade names or commercial products in this manuscript are solely for the purpose of providing specific information and do not imply recommendation or endorsement.

